# IOBRpy enables agentic multi-omics decoding of anti-tumor immunity

**DOI:** 10.64898/2026.07.17.739055

**Authors:** Haonan Huang, Xujia Li, Linfeng Liu, Wenchao Gu, Gaofeng Wang, Dongqiang Zeng

## Abstract

Decoding the tumor immunity is pivotal for cancer immunotherapy, yet transcriptomic pipelines remain bottlenecked by fragmented tools and biased interpretations. Here we present IOBRpy, a Python toolkit driven by an innovative AI dual-agent layer for automated, highly standardized immuno-oncology workflows. Moving beyond conventional expression profiling, IOBRpy enables agentic multi-omics decoding. From raw FASTQ or TPM matrices, it seamlessly integrates upstream quality control, transcript quantification, and downstream TME parsing, encompassing signature scoring, ligand-receptor crosstalk, and cellular deconvolution. Crucially, IOBRpy expands data dimensions by incorporating complementary immunogenomic layers, empowering concurrent high-resolution SpecHLA typing and TRUST4-based TCR/BCR repertoire reconstruction from sequencing data. Deployed across two large-scale cohorts (IMvigor210 and OAKPOPLAR), IOBRpy successfully captured multi-dimensional prognostic insights. While broad HLA-I heterozygosity showed negligible impact, it precisely unmasked treatment-stratified, allele-specific survival associations (e.g., HLA-A*01 and HLA-DPA1*02) tightly coupled with distinct immunosuppressive ligand-receptor networks (such as HMGB1-THBD and EFNB2-EPHB6) and dynamic TCR clonal diversity shifts. Empowering this lifecycle is a paired agent framework: a workflow agent automatically audits project states to execute validated, path-aware commands, while a result agent evaluates tool provenance, handles method-aware adaptive visualizations, and organizes findings into evidence-constrained biological hypotheses. Collectively, IOBRpy provides a reproducible, scalable, and intelligence-augmented Python gateway to transform raw sequencing data into multi-omic, interpretation-ready discoveries for cohort-scale precision immunotherapy (https://iobr.github.io/IOBRpy/).

## Introduction

The tumour microenvironment (TME) is an active cancer ecosystem in which malignant cells interact with immune and stromal populations, extracellular matrix, vasculature and soluble signalling networks. Rather than merely accompanying tumour growth, these components can drive tumour initiation, progression and metastatic dissemination and can shape malignant phenotypes independently of genetic change ^1-3^. Chemokine-guided cell trafficking, cancer-cell-intrinsic immunoregulatory programmes and suppressive myeloid populations organize the composition, spatial distribution and functional state of the immune milieu ^4-6^. This organization influences whether anti-tumour lymphocytes infiltrate and remain functional: T cell paralysis, variation in the abundance, state and localization of tumour-infiltrating lymphocytes, and immune selection of heterogeneous tumour subclones can promote immune escape and limit responses to immune-checkpoint blockade ^7-9^. The TME is therefore both a mechanistic determinant of cancer evolution and a clinically informative source of features for prognosis and treatment response ^1-9^.

This biological complexity creates a practical need for computational frameworks that convert transcriptomic data into interpretable TME readouts. Bulk RNA-seq remains widely available across public cohorts and translational studies, and established methods can infer immune infiltration, stromal abundance, tumour purity, pathway activity and immune-state signatures from expression profiles ^10-21^. Yet these methods are dispersed across programming languages, web services, scripts and incompatible input-output conventions. Starting from raw reads therefore requires users to connect preprocessing, expression quantification, matrix harmonisation, deconvolution, signature scoring, visualisation and interpretation. As workflows extend from expression-derived TME features to HLA genotypes and immune-repertoire outputs, users must also preserve data provenance, method-specific assumptions and valid comparison boundaries across heterogeneous evidence layers. The challenge therefore extends beyond fragmented code to non-standardised analysis and result over-interpretation: inappropriate transformations or comparisons can propagate through the workflow, while biologically plausible narratives can exceed the supporting evidence. Manually reconstructing and auditing this expanding analytical chain increases technical burden and weakens reproducibility.

To address this computational fragmentation, our group previously developed the IOBR R framework and IOBR 2.0, which have become highly cited and globally utilised benchmark resources for multi-omics TME profiling ^22,23^. This widespread academic recognition provided the empirical validation and practical rationale for IOBRpy: to extend this proven integrative philosophy to the Python ecosystem, where modern data-science and machine-learning workflows are increasingly anchored. IOBRpy does not alter the assumptions of the underlying TME methods. Instead, it provides a Python-native execution layer that improves access, workflow integration, reproducibility and extensibility.

A central design goal of IOBRpy is to connect upstream RNA-seq processing with downstream TME analysis while preserving the boundaries between distinct modules. Its FASTQ workflow combines read-level quality control, cohort reporting, Salmon- or STAR-based expression generation, matrix preparation and core TME profiling. HLA typing and immune-repertoire reconstruction remain complementary modules for aligned data ^24-31^. This organisation reduces manual transfer between tools while allowing users to select only the analyses supported by their inputs. IOBRpy is organised around project states and result types rather than a single monolithic pipeline. Its modules cover sequencing quality control, expression generation, HLA analysis, TCR/BCR reconstruction, immune and stromal deconvolution, tumour-purity estimation, signature scoring, clustering and ligand-receptor analysis ^10-21,24-31^. Agent-facing components add path-aware workflow guidance, result auditing, visualization and structured interpretation without changing the underlying analytical methods.

Here we describe the architecture and evaluation of IOBRpy. We first define the routes from raw sequencing data or expression matrices to TME profiling and complementary HLA/TCR modules. We then assess output concordance and runtime behaviour against corresponding R workflows for representative TME methods. Finally, we apply IOBRpy to two immuno-oncology cohorts and examine how paired agent interfaces support path-aware workflow selection, output auditing, adaptive visualization and evidence-constrained interpretation.

## Results

### IOBRpy connects raw RNA-seq processing to integrated TME analysis

IOBRpy provides entry points matched to common RNA-seq project states. Raw FASTQ data enter a workflow that performs quality control, Salmon- or STAR-based expression generation, matrix preparation, signature scoring, a default six-method TME panel and ligand-receptor scoring. Existing TPM matrices can enter directly at the downstream TME stage, whereas aligned BAM files are handled by separate HLA-typing and TRUST4 modules. MCP-backed agent interfaces expose path-mapping results, native commands and project state to command-aware assistants (Fig. 1a). At the module level, IOBRpy links fastp and MultiQC, Salmon or STAR, SpecHLA, TRUST4, signature scoring, immune deconvolution, clustering and ligand-receptor analysis (Fig. 1b) ^24^-^31^.

**Figure 1.**
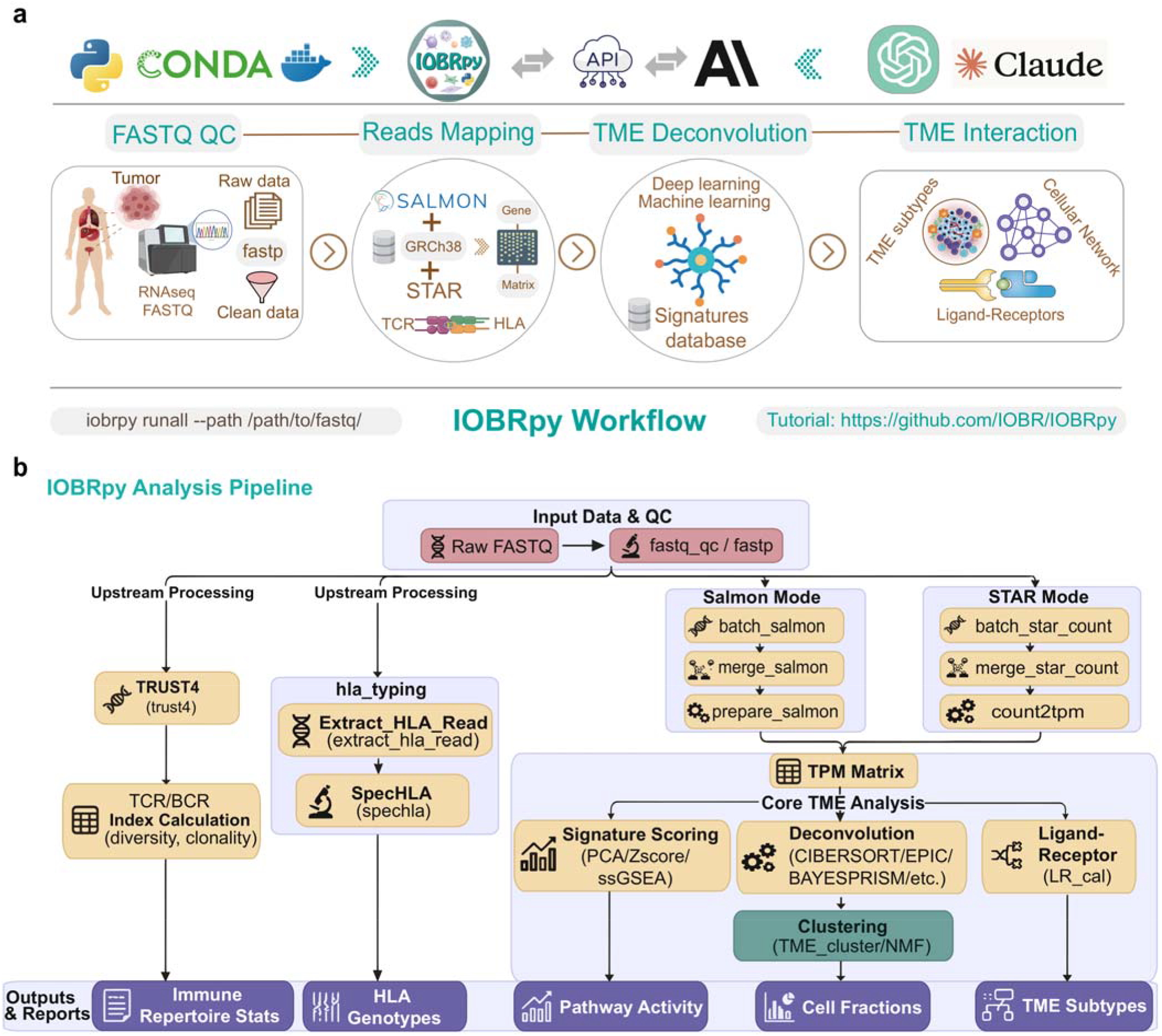
IOBRpy connects RNA-seq processing, TME profiling and immunogenomic analysis. (A) Conceptual architecture linking FASTQ quality control, read mapping, TME deconvolution and interaction analysis within a Python framework that supports Bioconda or container deployment and API/MCP-backed AI interfaces. (B) Modular routes from raw FASTQ data through fastp quality control, Salmon- or STAR-based expression generation, TRUST4 TCR/BCR reconstruction and SpecHLA typing. Downstream modules include signature scoring, the default six-method TME panel, separately available BayesPrism and clustering, and ligand-receptor analysis. Outputs include expression matrices, repertoire statistics, HLA genotypes, pathway activities, cell fractions and TME subtypes.

The complete workflow is organised around a standard project structure rather than a collection of independent command outputs. Upstream quality-control and expression-generation steps remain distinct from downstream TME modules, but their outputs are linked within the same analysis directory.A Resumable execution and top-level resource settings allow large cohorts to be processed incrementally while preserving a predictable relationship among inputs, intermediate matrices and downstream immune features.

The matrix-based workflow applies the same downstream logic without repeating FASTQ processing. From a TPM matrix, the default profiling route calculates signature scores, runs CIBERSORT, EPIC, quanTIseq, MCP-counter, ESTIMATE and IPS, merges their outputs by sample and calculates ligand-receptor scores. BayesPrism is provided as a separate deconvolution module with bundled single-cell expression data and matching cell-type and cell-state labels; users may instead supply a custom reference. Clustering is also run separately, keeping the default TPM workflow focused on the six-method panel and ligand-receptor analysis.

HLA and immune-repertoire modules extend expression-based profiling with complementary immunogenomic evidence. The batch HLA route extracts HLA-informative reads from aligned BAM files, applies SpecHLA and merges allele-level calls ^30^. TRUST4 accepts BAM or paired FASTQ inputs to reconstruct TCR/BCR sequences and summarise clonal richness, diversity, evenness and dominance ^31^. A single path-aware project configuration links the expression matrix and aligned sequencing data to TME profiling, HLA typing and TCR/BCR repertoire reconstruction, allowing the workflow-facing agent to coordinate all three dimensions without user-written glue code or manual path reconciliation.

### Python-native TME modules preserve R-derived outputs with method-specific runtimes

We next assessed whether Python-native execution preserved the behaviour of established TME methods. Representative IOBRpy modules were compared with corresponding R workflows for CIBERSORT, quanTIseq, MCP-counter, ESTIMATE, EPIC and IPS, spanning immune-cell deconvolution, stromal and tumour-purity estimation and immunophenotype-related scoring ^10-14,21^. We evaluated both runtime and output concordance because practical implementation requires computational tractability without loss of biological comparability.

Runtime profiles differed by method rather than by programming language alone (Fig. 2a). In the 936-sample benchmark, mean CIBERSORT runtime decreased from 2,864.8 s with one Python thread to 368.4 s with eight threads and 209.7 s with 16 threads, yielding a 13.7-fold speed-up from one to 16 threads. Python was also faster for quanTIseq, ESTIMATE and EPIC in this benchmark, whereas R showed shorter recorded runtimes for MCP-counter and IPS. The Python measurements included input-data loading, whereas R timing began after the data had been loaded; the apparent R advantage for some modules may therefore partly reflect the different timing scopes and should not be interpreted as a strict end-to-end comparison. The observed CIBERSORT scaling demonstrates parallel throughput at cohort scale. More broadly, Python’s in-memory reuse of loaded matrices, together with resumable module execution, persisted intermediates and configurable parallel resources, enables large cohorts to be partitioned and processed across distributed compute environments. These capabilities support TCGA- and ICGC-scale analyses and shift the relevant performance criterion from the runtime of an individual method to sustained throughput across large datasets.

**Figure 2.**
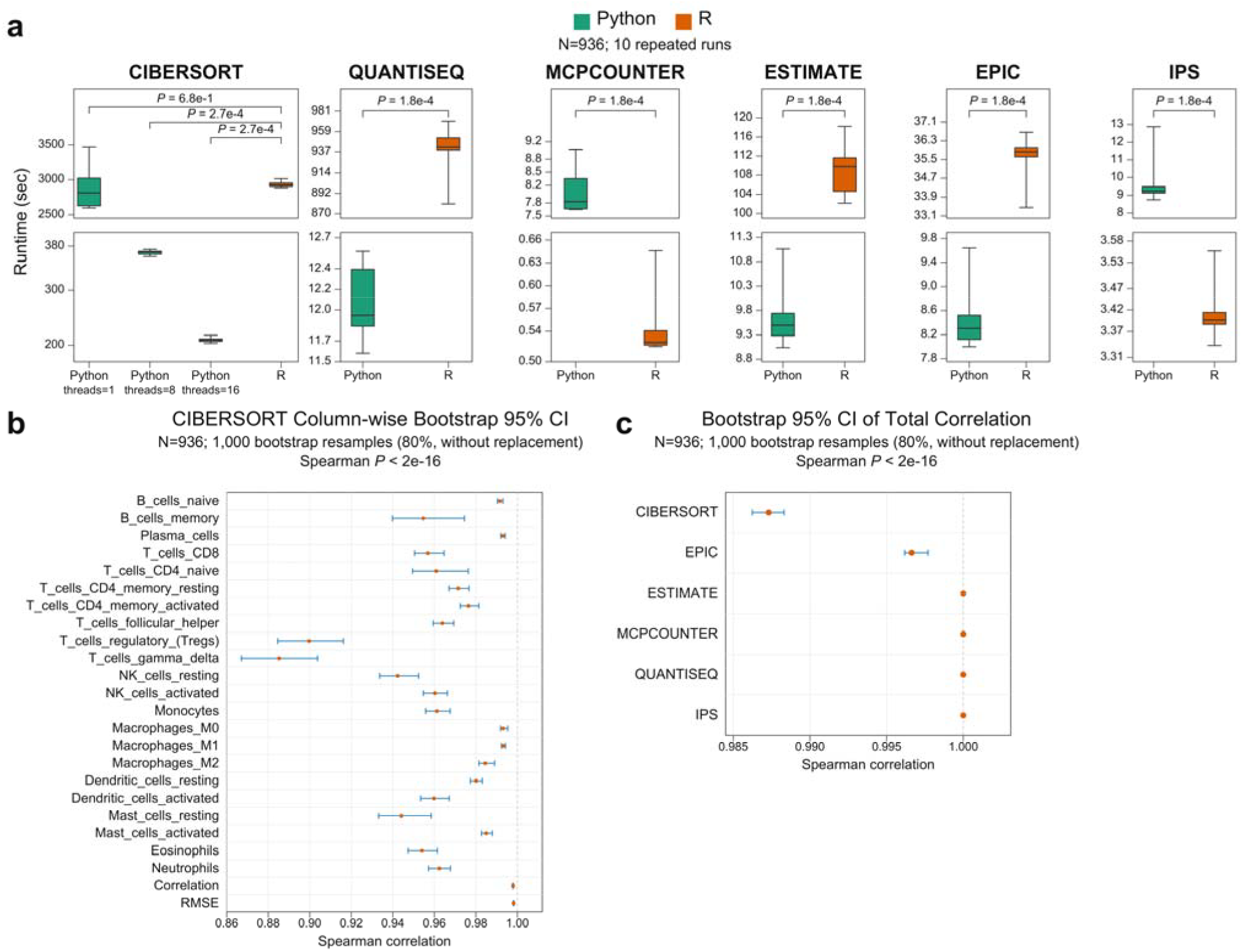
Runtime and output concordance of Python and R TME workflows. (A) Runtime distributions for CIBERSORT, quanTIseq, MCP-counter, ESTIMATE, EPIC and IPS. CIBERSORT includes Python runs using 1, 8 and 16 threads alongside the R workflow; the remaining panels compare Python and R implementations. Python timings included input-data loading, whereas R timings began after input loading. (B) Column-wise bootstrap 95% confidence intervals for CIBERSORT cell-type outputs. (C) Bootstrap 95% confidence intervals for total Spearman correlations between Python and R outputs.

Outputs were highly concordant across the tested modules (Fig. 2c). Total Spearman correlations between Python and R results were approximately 1.0 for IPS, quanTIseq, MCP-counter and ESTIMATE, 0.997 for EPIC and 0.986 for CIBERSORT. The lower aggregate correlation for CIBERSORT reflected cell-type-specific variation rather than global disagreement. Column-wise bootstrap confidence intervals further confirmed high concordance for most CIBERSORT fractions, with wider intervals for selected immune-cell subsets (Fig. 2b). Thus, IOBRpy retained the outputs of the tested TME algorithms while exhibiting module-specific computational profiles. Its principal value lies in reproducible workflow integration rather than a universal runtime advantage.

### Exploratory HLA/TCR analysis nominates treatment-stratified immune associations

We next asked whether IOBRpy could support integrated immunogenomic analysis beyond cell-fraction estimation. The case study used trial-derived data from IMvigor210, a multicentre single-arm phase II study of atezolizumab in advanced urothelial carcinoma (NCT02108652), and from the pooled OAK and POPLAR (OP) cohort, comprising the randomised phase III OAK trial (NCT02008227) and randomised phase II POPLAR trial (NCT01903993) in non-small-cell lung cancer ^32,33^. Within OP, immunotherapy (OP-IO) and chemotherapy (OP-Chemo) were treatment-defined subgroups rather than additional cohorts (Fig. 3a). SpecHLA-derived allele and zygosity features were analysed in both cohorts (Supplementary Table 2), and TRUST4-derived repertoire metrics were generated for both IMvigor210 and OP (Supplementary Table 4) ^30,31^. Treatment-stratified survival analysis of repertoire metrics was performed in OP.

**Figure 3.**
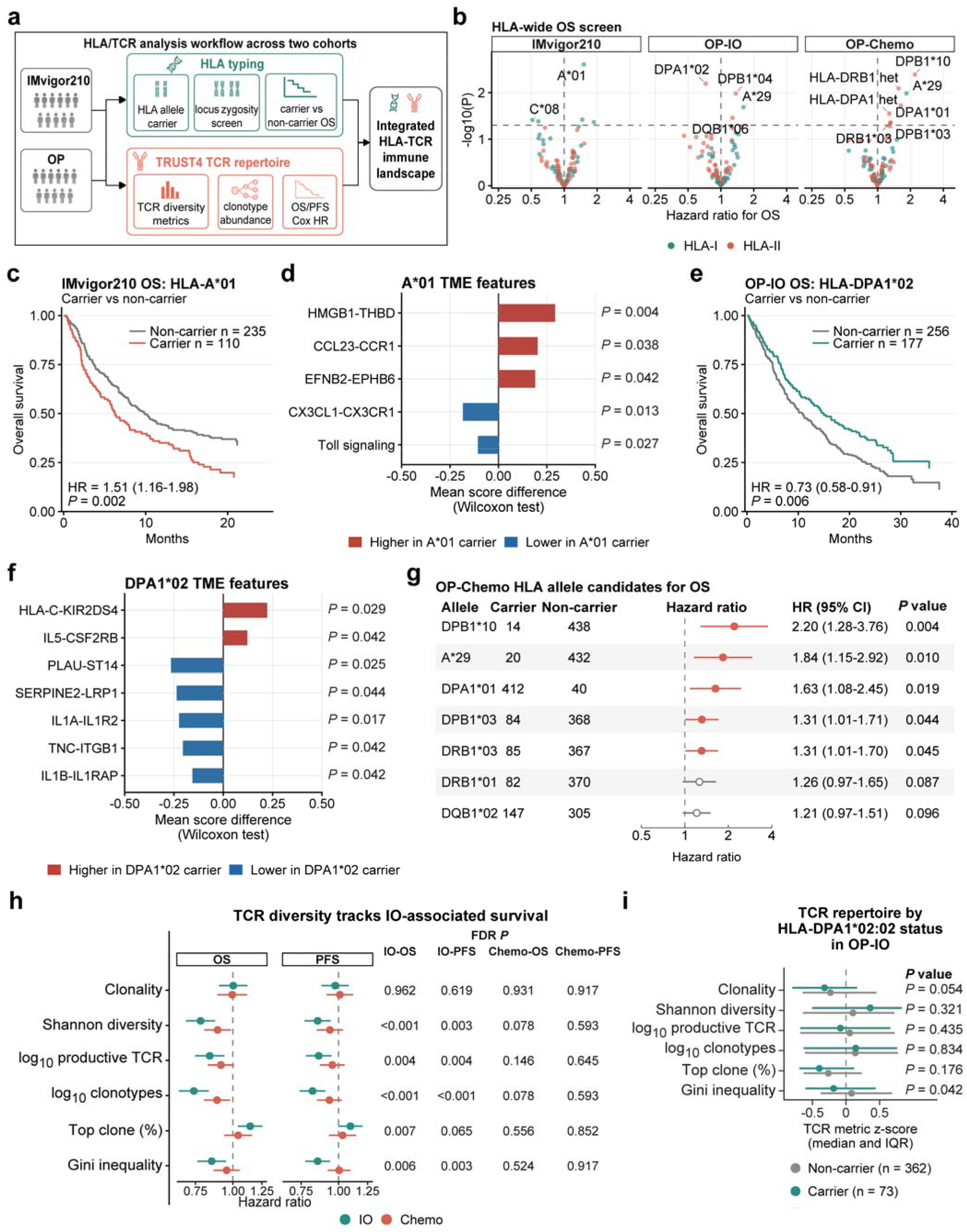
Integrated HLA/TCR and TME-feature analysis across two immuno-oncology cohorts. (A) Coordinated SpecHLA-based HLA and TRUST4-based TCR workflow across IMvigor210 and OP; OP was stratified as OP-IO and OP-Chemo. (B) HLA-wide screen of allele-carrier and locus-heterozygosity features for overall survival. (C) Kaplan-Meier curve for HLA-A*01 in IMvigor210. (D) Exploratory TME-associated features selected from IOBRpy-derived immune-deconvolution results, signature scores, ligand-receptor scores and repertoire metrics using IOBR-based Wilcoxon analysis (P < 0.05), comparing HLA-A*01 carriers with non-carriers in IMvigor210; Toll signaling is a signature score. (E) Kaplan-Meier curve for one-field HLA-DPA1*02 in OP-IO. (F) Exploratory TME-associated features selected using the same workflow, comparing one-field HLA-DPA1*02 carriers with non-carriers in OP-IO. (G) Selected OP-Chemo HLA allele candidates with at least 10 carriers and 10 non-carriers; HLA panels show nominal P values. (H) Hazard ratios per 1-s.d. increase for six TCR metrics across OS and PFS in OP-IO and OP-Chemo; FDR-adjusted P values are shown beside each endpoint. (I) Six TCR repertoire metrics compared between two-field HLA-DPA1*02:02 carriers (n = 73) and non-carriers (n = 362) in OP-IO. Points and horizontal lines indicate group medians and interquartile ranges of z-score-standardised metrics, respectively. Annotated P values were obtained from two-sided Wilcoxon rank-sum tests on original-scale values.

Broad HLA-I heterozygosity showed limited association with overall survival across IMvigor210, OP-IO and OP-Chemo: none of the all-locus, exactly two-locus, single-locus or pairwise definitions reached nominal significance (P < 0.05; Supplementary Table 1). We therefore used an HLA-wide screen to examine allele-carrier and locus-heterozygosity features across the three analysis groups (Fig. 3b). In IMvigor210, HLA-A*01 carriage showed a nominal association with shorter overall survival (110 carriers and 235 non-carriers; 230 events; HR = 1.51, 95% CI 1.16-1.98, P = 0.0024; Fig. 3c). In OP-IO, HLA-DPA1*02 carriage showed a nominal association with longer overall survival (177 carriers and 256 non-carriers; 312 events; HR = 0.73, 95% CI 0.58-0.91, P = 0.0064; Fig. 3e).

To place these exploratory allele-level associations in a TME context, we compared IOBRpy-derived TME features between the corresponding carrier and non-carrier groups using IOBR-based Wilcoxon analysis (P < 0.05). In IMvigor210, A*01 carriers showed higher ligand-receptor scores for HMGB1-THBD, CCL23-CCR1 and EFNB2-EPHB6, together with lower CX3CL1-CX3CR1 ligand-receptor scores and lower Toll signaling signature scores (Fig. 3d). EFNB2 encodes ephrin-B2, the ligand component of this pair. Representative supporting studies connect HMGB1, ephrin-B2 and CX3CL1/CX3CR1 with damage-associated signalling, urothelial immunotherapy resistance and T-cell-response biology ^34-36^. In OP-IO, DPA1*02 carriers showed higher ligand-receptor scores for HLA-C-KIR2DS4 and IL5-CSF2RB and lower scores for PLAU-ST14, SERPINE2-LRP1, IL1A-IL1R2, TNC-ITGB1 and IL1B-IL1RAP (Fig. 3f). IL5-CSF2RB was summarised as an IL-5/eosinophil-related signal because IL-5 is a canonical eosinophil-associated cytokine, and TNC-ITGB1 was summarised as tenascin-C because TNC encodes the extracellular-matrix protein tenascin-C. Representative supporting studies on IL-5/eosinophil biology, tenascin-C and IL-1β support immune-response, extracellular-matrix and inflammatory contexts ^37-39^. These exploratory feature differences provide biological context for the allele-level associations and nominate follow-up hypotheses.

Within OP-Chemo, the adequately represented candidate alleles selected for display in overall-survival analyses included DPB1*10, A*29, DPA1*01, DPB1*03 and DRB1*03, with hazard ratios above 1 (Fig. 3g). TCR repertoire analysis provided a complementary treatment-stratified layer (Fig. 3h). Clonality showed little association with either endpoint in either treatment group, but was retained as an evenness-derived summary of overall repertoire concentration. In OP-IO, five of six repertoire metrics for overall survival and four for progression-free survival reached the within-endpoint significance threshold (FDR-adjusted P < 0.05); corresponding chemotherapy-group associations did not pass the same threshold.

Given that TCR–pMHC interactions are a critical component of antitumor immunity and HLA molecules directly dictate T-cell antigen recognition specificity, to further investigate differences in TCR repertoire metrics between HLA-DPA1*02 carriers and non-carriers, we extended the analysis to two-field resolution and focused on HLA-DPA1*02:02 (OP-IO, 73 carriers and 362 non-carriers; Fig. 3i). Carriers showed a nominally lower Gini index and a concordant trend towards lower clonality, consistent with a more even, less dominance-skewed repertoire. Taken together, these analyses integrate HLA variation, survival associations, TME-associated features and TCR repertoire architecture across IMvigor210 and OP, illustrating how IOBRpy supports traceable exploratory immunogenomic analyses across treatment contexts.

### Paired agents separate workflow execution from result interpretation and visualisation

Finally, IOBRpy organises agent-supported analysis as a staged natural-language-guided workflow (Fig. 4). The workflow-facing agent interprets the request, assesses the available evidence and project state, ranks the next actions and asks the user to confirm the selected route. Guardrails then evaluate whether the proposed analysis is compatible with the available data and intended statistical design before local execution. They can block actions that would yield invalid or misleading results, such as testing associations across features on incompatible scales without the required z-score standardisation, or estimating TME features directly from expression data that have not undergone the required normalisation. By retaining the request, evidence, selected action and validation outcome, this stage provides a traceable transition from natural-language intent to a validated analysis.

**Figure 4.**
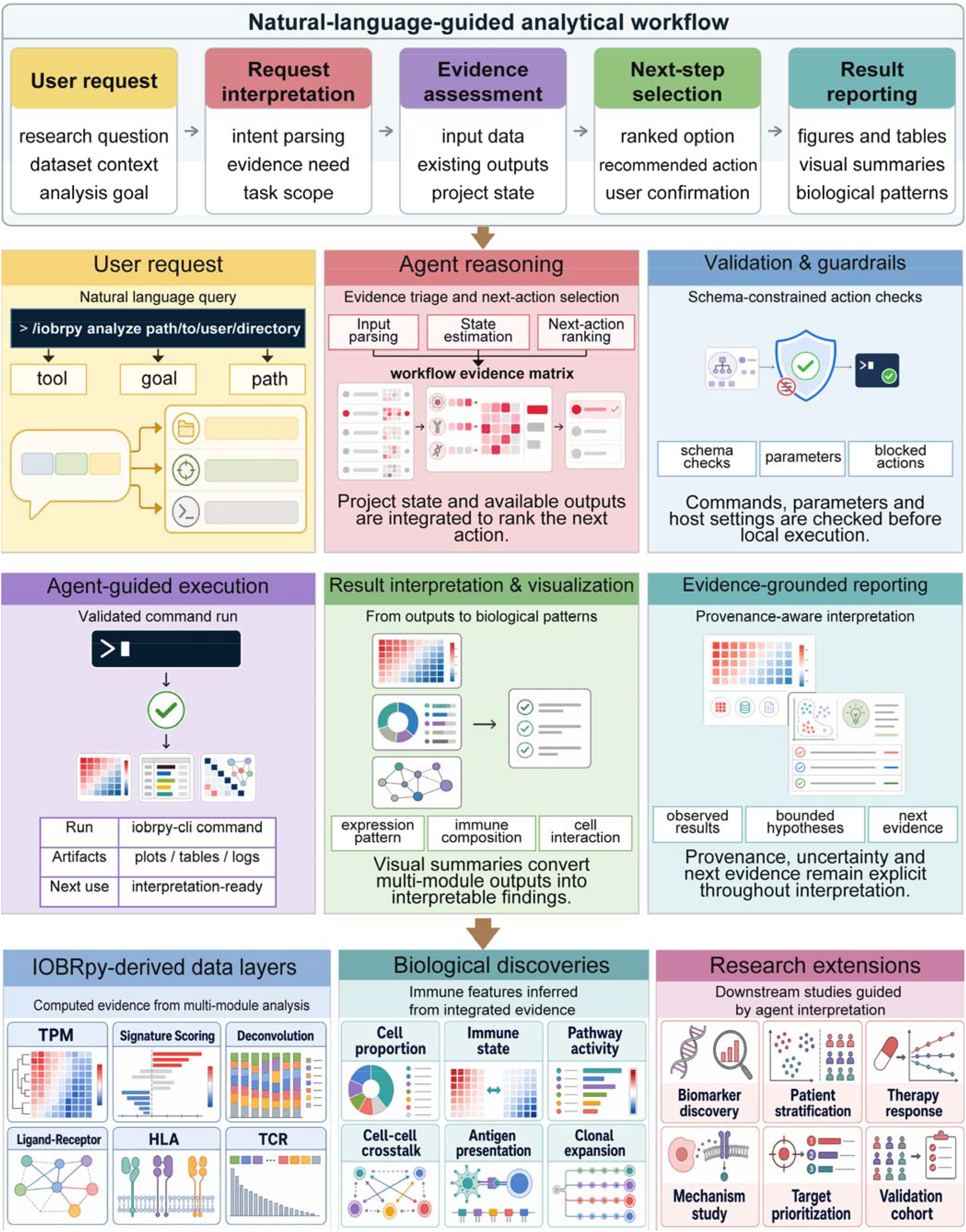
Natural-language-guided analytical workflow in IOBRpy. The upper sequence maps a user request through request interpretation, evidence assessment, ranked next-step selection and result reporting. The middle panels detail parsing of the requested tool, goal and path; evidence-based state estimation and action ranking; schema-constrained validation; execution of a validated IOBRpy command; result interpretation and visualisation; and provenance-aware reporting that separates observed results, bounded hypotheses and next evidence. The lower panels show how IOBRpy-derived data layers can be organised into candidate biological patterns and used to prioritise downstream biomarker, patient-stratification, mechanism, target and validation studies.

After execution, the result-facing agent converts validated outputs into figures, tables and visual summaries and organises them into evidence-grounded reports. Factuality constraints keep provenance and uncertainty explicit and separate observed results from bounded biological hypotheses and the next evidence required for validation. They prevent visual or narrative overstatement, for example by preserving the native scale of cell-composition estimates, avoiding claims of proven interactions from ligand-receptor co-expression scores, and keeping exploratory associations distinct from causal or clinical conclusions. Together, the paired agents connect validated execution with interpretable visualisation and evidence-bounded result interpretation.

### Simulated tasks demonstrate agent-guided analysis across TME, HLA and TCR outputs

To demonstrate the agent-facing workflow in a compact and traceable setting, we selected five OP-IO and five OP-Chemo samples and posed simulated natural-language tasks covering project inspection, analysis planning, workflow execution and result visualisation. The first panel summarises the sample set and the complementary roles of the workflow-facing and result-facing skills (Fig. 5a). A representative dialogue then shows path inspection, checklist-based analysis recommendation, execution of the selected workflow and hand-off of generated outputs for interpretation and visualisation (Fig. 5b).

**Figure 5.**
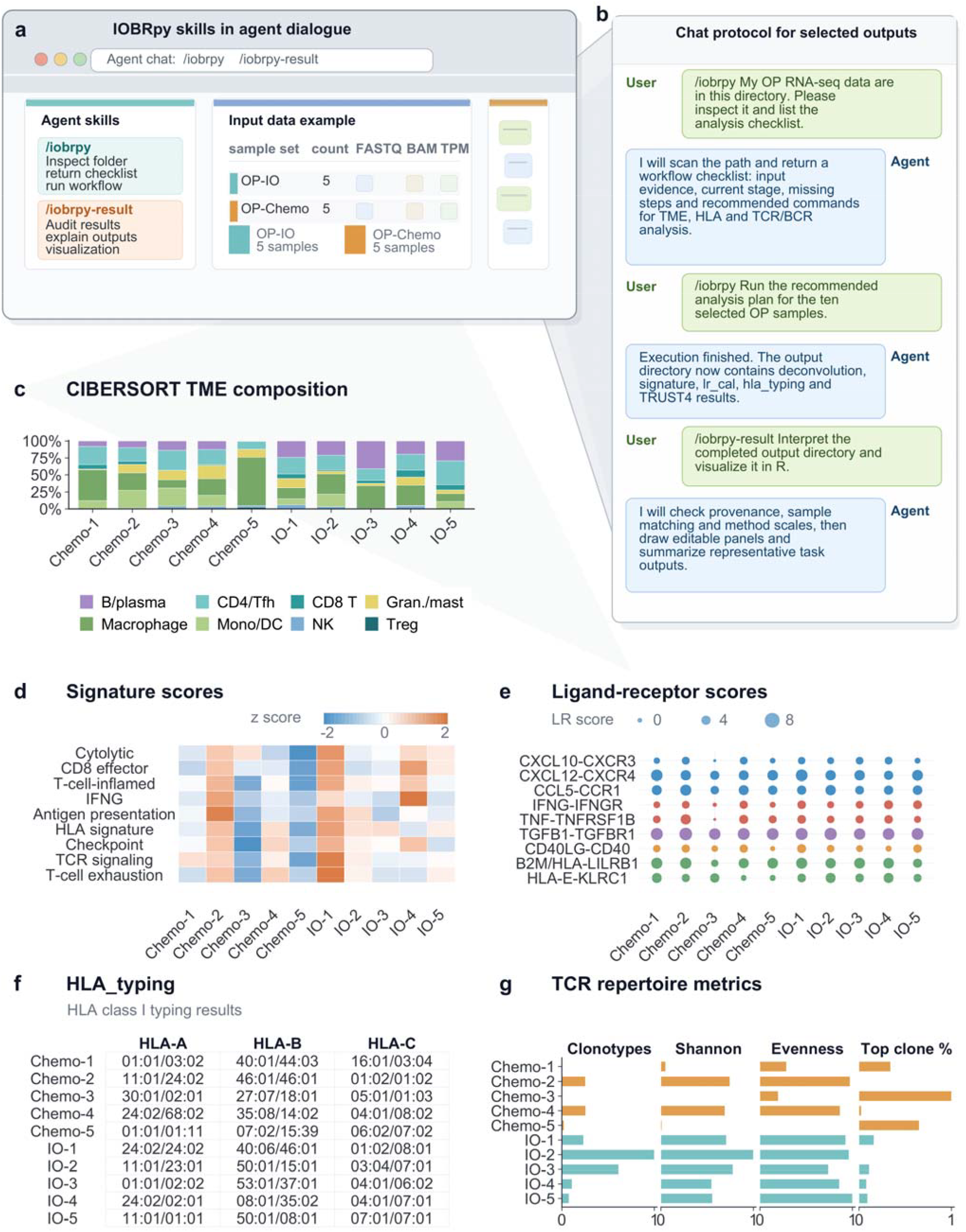
Ten-sample demonstration of agent-guided TME, HLA and TCR analysis. (A) Workflow-facing and result-facing skills with an input summary for five OP-IO and five OP-Chemo samples. (B) Representative dialogue covering path inspection, analysis recommendation, workflow execution and result visualisation. (C) CIBERSORT TME-composition visualisation. (D) Signature-score visualisation. (E) Ligand-receptor-score visualisation. (F) Tabulated HLA class I typing results. (G) TCR repertoire-metric visualisation.

The resulting outputs illustrate the data products handled within the same interaction sequence. CIBERSORT estimates summarise TME composition across the ten samples (Fig. 5c), while signature and ligand-receptor scores capture complementary immune-state and cell-communication features (Fig. 5d, e). Tabulated HLA class I calls extend the output to antigen-presentation features (Fig. 5f), and TCR metrics summarise clonal richness, diversity, evenness and dominance (Fig. 5g). These tasks connect natural-language project inspection and command execution to module-aware TME, HLA and TCR outputs while preserving the scale and structure of each result type.

## Discussion

IOBRpy addresses a practical source of fragmentation in immuno-oncology informatics. Informative TME methods are available, but moving from raw sequencing data or an expression matrix to integrated, reviewable outputs still requires substantial tool coordination. IOBRpy connects FASTQ-to-TME and TPM-to-TME workflows with separate HLA and repertoire modules, command-aware workflow guidance and result-facing interpretation rules. This integration reduces the engineering work required to move from RNA-seq inputs to coordinated TME and immunogenomic analyses.

The principal contribution of IOBRpy is an executable and inspectable integration layer rather than a new deconvolution model. Existing TME methods retain distinct assumptions, reference profiles and domains of applicability ^10-21^. IOBRpy preserves these method-specific properties while making the tools easier to combine, compare across cohorts and connect to upstream RNA-seq processing. It also exposes project state and generated outputs to command-aware agents. In this respect, IOBRpy extends rather than replaces the R-based IOBR framework ^22,23^.

The benchmark establishes technical concordance without implying a universal runtime advantage. Python and R outputs agreed closely for the tested modules, but computational behaviour varied by method and thread setting. Moreover, Python timing included input loading whereas R timing began after loading, so the recorded runtimes are descriptive rather than strictly equivalent end-to-end measurements. The supported conclusion is that IOBRpy standardises and connects the workflow while retaining module-specific computational trade-offs.

The HLA/TCR case study illustrates the value of combining immune evidence layers. HLA genotype and zygosity describe antigen-presentation potential, whereas TCR diversity and clonality capture complementary features of the adaptive immune response. Across the two cohorts and the OP treatment subgroups, broad HLA-I heterozygosity showed limited survival association, while candidate allele- and repertoire-level associations were observed in treatment-defined analyses. The Fig. 3d/F feature screens further placed these allele associations into TME contexts, but their feature-level biological interpretation remains hypothesis-generating; representative supporting studies for HMGB1, ephrin-B2, CX3CR1, IL-5/eosinophil, tenascin-C and IL-1β biology are cited in the main text ^34-39^. These findings demonstrate the analytical scope of the workflow and nominate candidate associations; they do not establish clinically actionable biomarkers.

The paired agent design addresses uncertainty at two different stages. Before or during analysis, the workflow-facing component scans project state, renders a complete checklist and recommends the next valid computational step. Native command schemas and Python guardrails constrain the selected action before execution. After analysis, the result-facing component classifies provenance, checks method-specific scales and transformation chains, supports adaptive visualisation and organises interpretation around observed results, bounded hypotheses and next evidence.

Several limitations define the current scope. IOBRpy inherits the assumptions and input requirements of its underlying methods, and estimates can be affected by sequencing depth, tissue processing, reference signatures and cohort composition. Bulk RNA-seq additionally conflates cell abundance, cell state and spatial organisation. Future versions will therefore combine single-cell references and spatial-transcriptomic measurements through multimodal large-model integration to better resolve these signals and reduce purity-related noise in bulk RNA-seq. Benchmark depth also differs among modules, warranting broader stress testing. The HLA/TCR associations and nominal TME-feature comparisons require independent validation before clinical or mechanistic interpretation. The result-facing agent depends on available metadata and companion files, and its recommendations, visual designs and structured interpretations should be quantified systematically across diverse result directories.

Future development will prioritise broader benchmarks, standardised reporting, stronger provenance capture and additional immune evidence layers. Planned directions include interfaces for single-cell and spatial transcriptomics, workflow-manager and cloud execution, expanded ligand-receptor resources and quantitative evaluation of agent-assisted command recommendation, visualisation success and interpretation factuality. Within its current scope, IOBRpy provides a Python-native framework for reproducible, inspectable and interpretation-ready TME analysis in translational immuno-oncology.

## Methods

### Software design and implementation

IOBRpy (v.0.2.0) is implemented as a Python command-line and workflow toolkit for RNA-seq-centred TME analysis. Analyses use Python (v.3.11) on Linux/POSIX systems, and the package is distributed through PyPI and Bioconda. Source code is available at https://github.com/IOBR/IOBRpy, and the online book and tutorials are available at https://iobr.github.io/IOBRpy/. The principal external workflow tools are fastp (v.1.3.6), MultiQC (v.1.31), Salmon (v.1.10.3), STAR (v.2.7.11b) and TRUST4 (v.1.1.9). HLA typing uses the SpecHLA implementation bundled with IOBRpy (v.0.2.0) and IPD-IMGT/HLA database (release 3.38.0). Supporting versioned dependencies include pysam (v.0.23.3), samtools (v.1.21), htslib (v.1.21), bcftools (v.1.21), freebayes (v.1.3.8), vcflib (v.1.0.10) and libdeflate (v.1.25). MCP-backed agent interfaces complement the native command surface with workflow guidance and result interpretation.

Commands are organised by analysis task. The runall workflow links FASTQ quality control, Salmon- or STAR-based expression generation, matrix preparation, signature scoring, the default six-method TME panel and ligand-receptor scoring. The tme_profile workflow starts from a TPM matrix and runs the same downstream TME stack. BayesPrism, clustering, HLA typing and TRUST4 remain separate modules because they use distinct inputs or outputs. Existing outputs can be reused through selective module execution or resume-aware workflow runs.

### FASTQ preprocessing and quality control

For raw RNA-seq input, IOBRpy performs read preprocessing and quality control with fastp (v.1.3.6) ^24-26^. Sample-level outputs are aggregated with MultiQC (v.1.31) to summarise read quality, filtering behaviour and sequencing consistency across the cohort ^27^. This quality-control layer provides a common upstream evidence base for both quantification- and alignment-based expression-generation routes.

### Expression generation and matrix preparation

IOBRpy supports Salmon- and STAR-based expression generation. In Salmon mode, per-sample quantifications are merged and converted to a cleaned gene-level TPM matrix ^28^. In STAR mode, aligned read-count outputs are merged and converted to TPM values ^29^. Both routes retain the untransformed TPM output and a log2(x+1) expression matrix in a standard directory structure for downstream analysis.

### HLA typing and TCR/BCR repertoire analysis

The batch HLA module scans a BAM directory, extracts HLA-informative reads and performs high-resolution typing with the SpecHLA implementation bundled in IOBRpy (v.0.2.0) against IPD-IMGT/HLA (release 3.38.0) ^30^. It returns per-sample results and a merged HLA genotype table. The TRUST4 (v.1.1.9) wrapper accepts individual or batched BAM and paired FASTQ inputs for TCR/BCR reconstruction ^31^. After reconstruction, IOBRpy writes clonotype-level data and sample-level summaries of richness, diversity, evenness and clonal dominance. Because aligned BAM files can support both SpecHLA and TRUST4 while matched expression matrices feed the TME modules, one path-aware project specification can coordinate TME, HLA and TCR/BCR analyses without user-written glue code. Separate native commands and method-specific input requirements are retained for reproducibility.

### TME deconvolution and signature scoring

The default TME panel comprises CIBERSORT, EPIC, quanTIseq, MCP-counter, ESTIMATE and IPS ^10-14,21^. Their outputs are merged by sample in the runall and tme_profile workflows. BayesPrism is provided as a standalone Bayesian deconvolution module that jointly models bulk expression and a single-cell reference ^15^. IOBRpy supplies a bundled single-cell expression reference with matching cell-type and cell-state labels, while allowing users to provide their own reference. BayesPrism remains outside the default panel because its inputs and output structure differ from those of the six bundled methods. Sample-level outputs from the six default methods and BayesPrism are provided for the IMvigor210 and OP case-study cohorts (Supplementary Table 3).

Signature scoring supports PCA, z-score, ssGSEA and integration modes ^16-20,40^. The tme_cluster and NMF modules operate separately on selected feature matrices to define sample groups. Ligand-receptor analysis calculates expression-derived interaction scores using user-specified data type, gene identifier type and cancer context.

### Benchmarking against R workflows

Python and R implementations of CIBERSORT, quanTIseq, MCP-counter, ESTIMATE, EPIC and IPS were benchmarked using the same 936-sample OP expression matrix and computing environment. Runtime was recorded across 10 repeated runs for each setting; Python timing included input-data loading, whereas R timing began after the data had been loaded. CIBERSORT was additionally evaluated with 1, 8 and 16 Python threads. Output concordance was quantified using Spearman correlation, with total correlations summarising module-level agreement and column-wise bootstrap 95% confidence intervals summarising CIBERSORT cell-type outputs. Bootstrap analysis used 1,000 draws of 80% of samples without replacement.

### HLA/TCR cohort analysis

The HLA/TCR case study comprised trial-derived data from IMvigor210, a multicentre single-arm phase II study in advanced urothelial carcinoma (NCT02108652), and the pooled OAK and POPLAR (OP) cohort from randomised phase III OAK (NCT02008227) and randomised phase II POPLAR (NCT01903993) trials in non-small-cell lung cancer ^32,33^. OP-IO and OP-Chemo were treatment-defined subgroups of OP rather than independent cohorts. The current biomarker analyses were retrospective and exploratory; the phase and randomisation of the parent trials do not confer prospective validation on the reported HLA or TCR associations. HLA features were analysed in both cohorts, whereas TCR repertoire features entered the survival analysis only in OP. Original full-resolution SpecHLA calls are provided for 348 IMvigor210 and 936 OP samples (Supplementary Table 2). For carrier and zygosity analyses in Fig. 3b–g, these calls were normalised to one-field allele notation. The direct HLA-TCR comparison in Fig. 3i retained two-field DPA1 calls to define HLA-DPA1*02:02 carriage. Samples with incomplete typing at a tested locus were excluded from that locus-specific comparison. HLA-wide screening tested allele-carrier status and locus heterozygosity against overall survival. Broad HLA-I definitions included full HLA-A/B/C heterozygosity, exactly two heterozygous loci, single-locus heterozygosity and pairwise HLA-A/B/C combinations (Supplementary Table 1).

Productive TCR clonotypes were identified from TRUST4 records with valid standard-amino-acid CDR3 sequences and TCR gene annotation and were deduplicated by chain, V gene, J gene and CDR3 amino-acid sequence. Six repertoire metrics were evaluated: clonality (1 - evenness), Shannon diversity, log10 productive TCR clonotype count, log10 total clonotype count, top-clone percentage and Gini inequality. The metrics were selected to represent complementary aspects of repertoire architecture rather than on the basis of survival significance: breadth (productive and total clonotype counts), overall diversity (Shannon diversity) and clonal abundance structure (clonality, top-clone percentage and Gini inequality). Clonality was retained because it is a standard distribution-wide measure derived as 1 - evenness and complements the richness and clonal-dominance metrics. Sample-level TRUST4-derived summary metrics are reported for 347 IMvigor210 and 936 OP samples (Supplementary Table 4); only OP samples entered the treatment-stratified survival models. Metrics were z-score standardised across the combined OP samples before treatment-stratified analyses. Accordingly, hazard ratios for continuous TCR metrics are reported per 1-s.d. increase. Univariate Cox models used Breslow handling for HLA and Efron handling for continuous TCR metrics; TCR P values within each endpoint-by-treatment set of six tests were adjusted using the Benjamini-Hochberg procedure and reported as FDR-adjusted P values. Kaplan-Meier curves visualised selected allele-level associations. The OP-Chemo forest panel required at least 10 carriers and 10 non-carriers for display. For the direct HLA-TCR comparison, 435 OP-IO samples with complete two-field HLA-DPA1 typing and all six TCR metrics were classified as HLA-DPA1*02:02 carriers (n = 73) or non-carriers (n = 362). Each metric was separately z-score standardised across these samples for visualisation, with group medians and interquartile ranges displayed. Two-sided Wilcoxon rank-sum tests were performed on original-scale values, with the six resulting raw P values reported. Exploratory TME-feature interpretation for the HLA-A*01 and HLA-DPA1*02 carrier comparisons used IOBRpy-derived outputs generated by cibersort, epic, ips, estimate, quantiseq, mcpcounter and bayesprism for immune deconvolution; calculate_sig_score for signature scoring; and lr_cal for ligand-receptor analysis, with TRUST4-derived TCR repertoire metrics included where available. Carrier and non-carrier groups were compared with the IOBR batch_wilcoxon function, and displayed features were selected by nominal Wilcoxon testing (P < 0.05).

### Workflow-facing natural-language analysis layer

The workflow-facing layer begins by parsing a natural-language request into the requested tool, analysis goal and data path. It then scans the user-specified directory, classifies detected evidence by workflow stage and renders a grouped checklist that retains both completed and missing items. The scan distinguishes IOBRpy-confirmed outputs from inferred or external results when provenance is uncertain. Native commands are exposed as structured MCP tools generated from the command parser. A language model proposes a structured next action from the evidence matrix, while packaged schemas and Python guardrails validate the command name, remove unknown parameter keys, check allowed choices, host-sensitive settings, input types and required files, and block local execution when mandatory evidence is missing. Command-specific rules prevent inappropriate handoffs, including raw FASTQ or directory inputs to TPM-only downstream commands and automatic reapplication of log transformation to an already transformed matrix.

### Result interpretation and visualisation layer

The result-facing layer operates on outputs that have already been generated. It first builds an evidence inventory, classifies candidate files by provenance and records their detected role, source class, decisive evidence and confidence. It then audits delimiter, table orientation, sample identifiers, duplicate or lost joins, numeric ranges, missingness, non-finite values and available uncertainty fields before constructing a result profile covering method family, unit, orientation, sample key, feature family, implementation variant and effective transformation chain. Files with ambiguous provenance or transformation status remain explicitly separated from confirmed native outputs and are not promoted to biological evidence.

Interpretation-only requests proceed from evidence classification to a structured report of identified results, field meanings, direct observations, method-aware interpretation, necessary limitations and next evidence. Figure requests require the user to select Python or R before plotting; the selected backend is then used for design, rendering, source-data export and visual quality control. The agent does not silently normalise, impute, filter or drop samples. Any required z-score transformation or other standardisation is made explicit and retained in the analysis record. Factuality rules preserve the biological meaning and scale of each result: fractions are not converted into absolute cell counts, relative abundance is not described as absolute abundance, ligand-receptor co-expression scores are not treated as proven interactions, and exploratory associations are not promoted to causal or clinically actionable claims.

### Statistical analysis

Runtime distributions were summarised across repeated runs using boxplots, with significance annotations shown in the figure. Output concordance was quantified by Spearman correlation. Survival analyses used Cox proportional-hazards models and Kaplan-Meier estimates. Adjusted significance thresholds were applied where multiple related features were screened.

## Data and code availability

IOBRpy source code is available at https://github.com/IOBR/IOBRpy. The analyses reported here used IOBRpy (v.0.2.0).

The IMvigor210 data used in the HLA/TCR case study are available through EGA study EGAS00001002556, including RNA-seq dataset EGAD00001003977. Data for the pooled OAK and POPLAR (OP) cohort are available through EGA study EGAS00001005013, including dataset EGAD00001007703. Access is controlled by the original data providers and may require application, approval and a data-use agreement; individual-level controlled-access data should not be redistributed. OP-IO and OP-Chemo are treatment-defined subgroups of the pooled OP cohort, not independent cohorts. No new patient samples were generated for this software study unless otherwise stated. Derived statistical summaries, original HLA calls, immune-deconvolution and scoring outputs, and TCR repertoire metrics supporting the case study are provided separately (Supplementary Tables 1-4).

## Acknowledgements

The work was supported by the National Natural Science Foundation of China (No. 82404016) and the Guangdong Basic and Applied Basic Research Foundation (No. 2023A1515110193) of D. Z. All funders had no involvement in any aspect of the study, including its design, collection, analysis, interpretation of the data, or the decision to submit the paper for publication.

## Author contributions

H.H. conceived the study with D.Z., designed and developed the complete IOBRpy software and paired agent framework, established the methodology, curated the data, performed the computational analyses and benchmarks, generated the figures, and wrote the original draft (Conceptualization, Methodology, Software, Validation, Formal analysis, Investigation, Data curation, Visualization, Writing - original draft, and Project administration). X.L. curated clinical metadata, validated cohort annotations and reviewed the manuscript (Data curation, Validation, and Writing - review and editing). L.L. independently checked statistical analyses and figure source data and reviewed the manuscript (Validation, Formal analysis, and Writing - review and editing). W.G. advised on the agent methodology, evaluated agent-facing workflows and reviewed the manuscript (Methodology, Validation, and Writing - review and editing). D.Z. co-conceived and supervised the study, provided resources and clinical interpretation, acquired funding and revised the manuscript (Conceptualization, Resources, Supervision, Funding acquisition, and Writing - review and editing). All authors reviewed and approved the manuscript.

## Supplementary table legends

Supplementary Table 1. Associations of HLA-I heterozygosity features with overall survival in IMvigor210, OP-IO and OP-Chemo.

Values are from univariate Cox models. Samples with incomplete typing at a required HLA locus were excluded from the corresponding comparison. Nominal P values and FDR-adjusted P values are shown (P < 0.05 or FDR-adjusted P < 0.05).

Supplementary Table 2. Sample-level full-resolution HLA calls in IMvigor210 and OP.

Original SpecHLA allele strings are provided for HLA-A, HLA-B, HLA-C, HLA-DPA1, HLA-DPB1, HLA-DQA1, HLA-DQB1 and HLA-DRB1 in 348 IMvigor210 and 936 OP samples. One-field transformations and derived carrier or zygosity variables used in downstream analyses are not included.

Supplementary Table 3. Sample-level immune-deconvolution and scoring outputs in IMvigor210 and OP.

Sample-level immune-deconvolution and scoring outputs from CIBERSORT, EPIC, ESTIMATE, IPS, MCP-counter, quanTIseq and BayesPrism are provided for 348 IMvigor210 and 936 OP samples.

Supplementary Table 4. Sample-level TRUST4-derived TCR repertoire summary metrics in IMvigor210 and OP.

Metrics are provided for 347 IMvigor210 and 936 OP samples; only OP samples were used in the treatment-stratified survival analysis shown in Fig. 3h.

